# Accurate estimation of canine inbreeding using ultra low-coverage whole genome sequencing

**DOI:** 10.64898/2026.04.04.716453

**Authors:** Ryan Kim, Declan Smith, Liudmilla Rubbi, Gregory Kislik, Matteo Pellegrini

**Affiliations:** Molecular Cell and Developmental Biology, University of California, Los Angeles (UCLA), Los Angeles, CA, United States; University of Washington, Seattle, WA, United States

## Abstract

The measurement of inbreeding has gained significance across diverse fields, including population and conservation genetics, agricultural genetics, breeding programs for animals and plants, and wildlife management. This is due to the fact that inbreeding leads to increased homozygosity and results in lower genetic diversity, rendering populations more vulnerable to environmental changes, diseases, and other stressors. High or mid-coverage whole genome sequencing (WGS) has been widely used for inbreeding estimation, but it is resource-intensive. We aimed to investigate the use of ultra low-coverage whole genome sequencing (ulcWGS) as a cost-effective alternative for inbreeding analysis. Domestic dogs were used for our study as their extensive breeding histories lead to populations with a wide range of inbreeding levels. We constructed a multi-breed reference panel from high-coverage WGS samples. Inbreeding in independent ulcWGS samples was then estimated using runs of homozygosity (RoH) and inbreeding coefficients (*F*). We modeled the relationship between these measures and sequencing depth using nonlinear regression, to generate inbreeding estimates relative to sequencing depth. Resulting relative RoH and F measurements were significantly correlated, with purebred dogs exhibiting more runs of homozygosity and higher inbreeding coefficients compared to mixed-breed dogs. Our findings demonstrate that ulcWGS can provide reliable and economical estimations of inbreeding, expanding accessibility to genetic monitoring.

## Introduction

Inbreeding refers to the mating of two individuals that share lineage from one or more common ancestors. All living organisms exhibit inbreeding to some extent, the levels of which can be evaluated through pedigree records or estimated via genomic data. Inbreeding leads to increased homozygosity and decreased heterozygosity. This effect is pronounced in small and isolated populations such as selective animal and plant breeding pools [1, 2]. The genetic consequences that arise from this phenomenon pose significant risks. First, allele diversity of the gene pool is reduced, lowering fitness of the population. This can make populations more vulnerable to environmental changes, diseases, and other stressors as reduced genetic diversity limits a population’s ability to adapt to changing environments or new challenges. Second, the accumulation of deleterious mutations increases the chances of their expression, reducing the individual’s fitness [3]. These negative effects, known as inbreeding depression, negatively influence reproduction and survival rates [4].

The practice of estimating inbreeding in both individuals and populations has become increasingly important in both agricultural and conservation genetics. However, pedigree information for most of the study subjects is often incomplete or inaccurate, necessitating the use of genomic analysis to infer inbreeding levels. Determining inbreeding levels accurately and cost-effectively through genomic data is of value for assessing health and fitness in individuals, managing livestock, and conserving endangered species [5], as for example Scandinavian wolves [6].

Thus far, WGS has been extensively used to estimate levels of inbreeding. With coverage higher than 15x, WGS enables the estimation of genetic variants across most of the genome. Whole-genome sequencing (WGS) enables high-resolution estimation of inbreeding by directly capturing genome-wide patterns of homozygosity and shared ancestry. One widely used approach is the identification of runs of homozygosity (ROH), long contiguous stretches of homozygous genotypes that arise from inheritance of identical chromosomal segments from recent common ancestors, summarized as the fraction of the genome in ROH, which provides a robust and interpretable measure of autozygosity [7, 8]. Complementary methods estimate inbreeding coefficients (F) based on the excess of observed homozygosity relative to Hardy– Weinberg expectations across genome-wide variant data, or via genomic relationship matrices that quantify identity-by-descent using allele frequency–weighted covariance among individuals [9-11]. Together, these WGS-based estimators capture distinct aspects of inbreeding history, with ROH-based measures being particularly sensitive to recent consanguinity, while homozygosity- and relationship-based coefficients integrate information across both recent and more ancient demographic processes.

While WGS is ideal for studies of inbreeding, its cost can be prohibitive for large projects. High-coverage WGS also produces large data volumes, which require significant compute resources. By contrast, lcWGS is more cost-effective and efficient, but limits the accuracy of genotype inferences [12].

Several studies have examined the use of low-coverage whole genome sequencing for genotyping and downstream analyses. One such study collected lcWGS data of rainbow trouts and found that a reference haplotype panel can be used to accurately impute the genotype [13]. Another study experimented with genotype imputation to estimate identity by descent tracts from low-coverage NGS data of rice [14]. Another study investigated the genomic diversity of North American grey wolves using lcWGS with mean depth of 7x [15]. However, the effectiveness of ultra low-coverage sequencing to assess inbreeding remains unclear [16].

In this study, we investigate the use of ultra low-coverage whole genome sequencing (0.1x-0.6x) for estimating inbreeding levels. We chose to examine canines because of their widespread use as a model for genetics [17]. Domestic canines are a useful species for studying inbreeding and genetic heredity owing to the history of descent from a population of wolves [18] and numerous population bottlenecks, along with social and artificial selection. In addition, numerous prior studies that have examined inbreeding in dogs using arrays and high-coverage WGS provides a useful reference for inbreeding research [19].

We computed inbreeding coefficients and runs of homozygosity in ultra low-coverage WGS data (< 1x) derived from 96 dogs. Our inbreeding estimates were corrected for read depth and a significant association between the two inbreeding metrics was found. Finally, we measured the relationship between RoH and F as a function of sequencing depth by downsampling one individual. We found that using our depth corrected metrics purebred dogs had higher inbreeding levels compared to mixed dogs, supporting the notion that low-pass genome sequencing with mean depth of 0.1x-0.6x can be used for studying inbreeding.

## Methods

### Whole genome sequencing

We collected 96 buccal swabs from dogs. For each sample, the following details were collected: date of birth, breed, weight, sex, and spayed/neutered status. Samples from these dogs were collected with the owners’ signed consents. The samples were collected from dog parks, breeders, and online volunteers (Princeton IACUC approved protocol # 2098A).

Buccal swab samples were collected using the PERFORMAgene kit (PG-100) from DNAGenotek, and DNA was isolated according to the manufacturer’s instructions. Samples were stored at −80 °C. DNA was extracted from the buccal swabs using the vendor-supplied protocol and incubated overnight at 50 °C before DNA extraction.

100 ng of extracted DNA was used for whole genome sequencing (WGS) library preparation. Fragmented DNA was subject to end repair, dA-tailing, and adapter ligation using the NEBNext Ultra II Library prep kit Library. QC was performed using the High-Sensitivity D1000 Assay on a 2200 Agilent TapeStation. Pools of 96 libraries were sequenced on a NovaSeq X Plus (10B lane) as paired-end 150 bases.

### Construction of reference population

To estimate expected allele frequencies in a population of dogs we utilized a set of 170 dogs from diverse breeds with high-coverage WGS. Fastq files were retrieved from the Sequence Read Archive (SRA, the full list of accession numbers and breeds can be found in Table S1) and aligned to the CanFam4 assembly with BWA-mem2 (version 2.2.1) [20]. Variants were called with bcftools (version 1.18) mpileup and call, then converted to binary files with PLINK 1.9 (www.cog-genomics.org/plink/1.9/) [10]. We genotyped a pool of ∼20 million linkage disequilibrium filtered variants from Meadows et al. (2023) [20]. These binary files were merged into a reference genome with PLINK -bmerge. Only biallelic variants on autosomal chromosomes were retained. An allele frequency report of these reference SNPs was generated using PLINK –freq. These reference allele frequencies serve as a basis for unbiased inbreeding coefficient calculations.

### Computing inbreeding metrics with PLINK

We computed inbreeding metrics using PLINK. To calculate the coefficient of inbreeding, PLINK 1.9’s –het command was used. PLINK calculates the expected autosomal heterozygous genotypes for each sample and reports a method-of-moments F estimate.

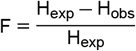

where H_exp_ is the expected heterozygous count and H_obs_ is the observed heterozygous count. The expected heterozygous count relies on accurate minor allele frequencies. Because calculating allele frequencies directly from our dataset could introduce bias, we utilized the allele frequency report of the reference population to calculate an unbiased expected heterozygous count for each sample using PLINK 1.9’s –read-freq flag.

To calculate the runs of homozygosity, we used PLINK 1.9’s –homozyg command. We set the report threshold to runs containing 100 SNPs that were longer than 1000 kilobases. This outputs the number of segments of RoH, the total kilobases, and the average kilobases.

### Modeling the inbreeding relative to coverage

We calculated the average coverage depth of each sample. Since coverage affects the calculated inbreeding metrics, we used LOESS regression to generate trends for F and RoH versus coverage. LOESS allows us to account for nonlinearities between inbreeding and coverage, as the estimations will converge at high coverage. This step created a reference inbreeding coefficient trend as a function of coverage. The inbreeding estimates corrected for coverage were obtained through the residuals of the actual values from those predicted by the reference model. High values of these residuals indicate higher inbreeding than expected

### Regression of inbreeding metrics against sequencing depth in synthetic data

The effect of sequencing depth on RoH and F values was mapped using downsampled versions of a single sample. One high-coverage WGS individual was selected. We used SAMtools –view to randomly select reads from a range of 1% to 39%, which best mimics the coverage range of the samples. The dataset was then processed through the same pipeline as the other samples, and the trend of depth to RoH and F was compared to that in the original dataset.

## Results

### Calculation of inbreeding metrics

We sequenced DNA collected from the buccal swabs of 96 dogs of varied breeds using low coverage whole genome sequencing (ulcWGS) with coverage around 0.1 to 0.5x. To obtain the expected allele frequencies across a panel of SNPs that had previously been reported from Meadows et al. (2023) [20], we downloaded a set of 170 dogs from SRA with varied breeds that had been sequenced at higher coverage (5-10x). For both sets of samples, the reads were aligned to the CanFam4 genome, and VCF files were generated using the bcftools mpileup and call commands (see Methods).

RoH and F values for each sample among the 96 ulcWGS dogs were calculated using PLINK (see Methods). The two metrics had a negative correlation with read depth (Fig 1). This is expected since lower sequencing depths will lead to overestimation of homozygous calls (Fig 1).

**Fig 1.**
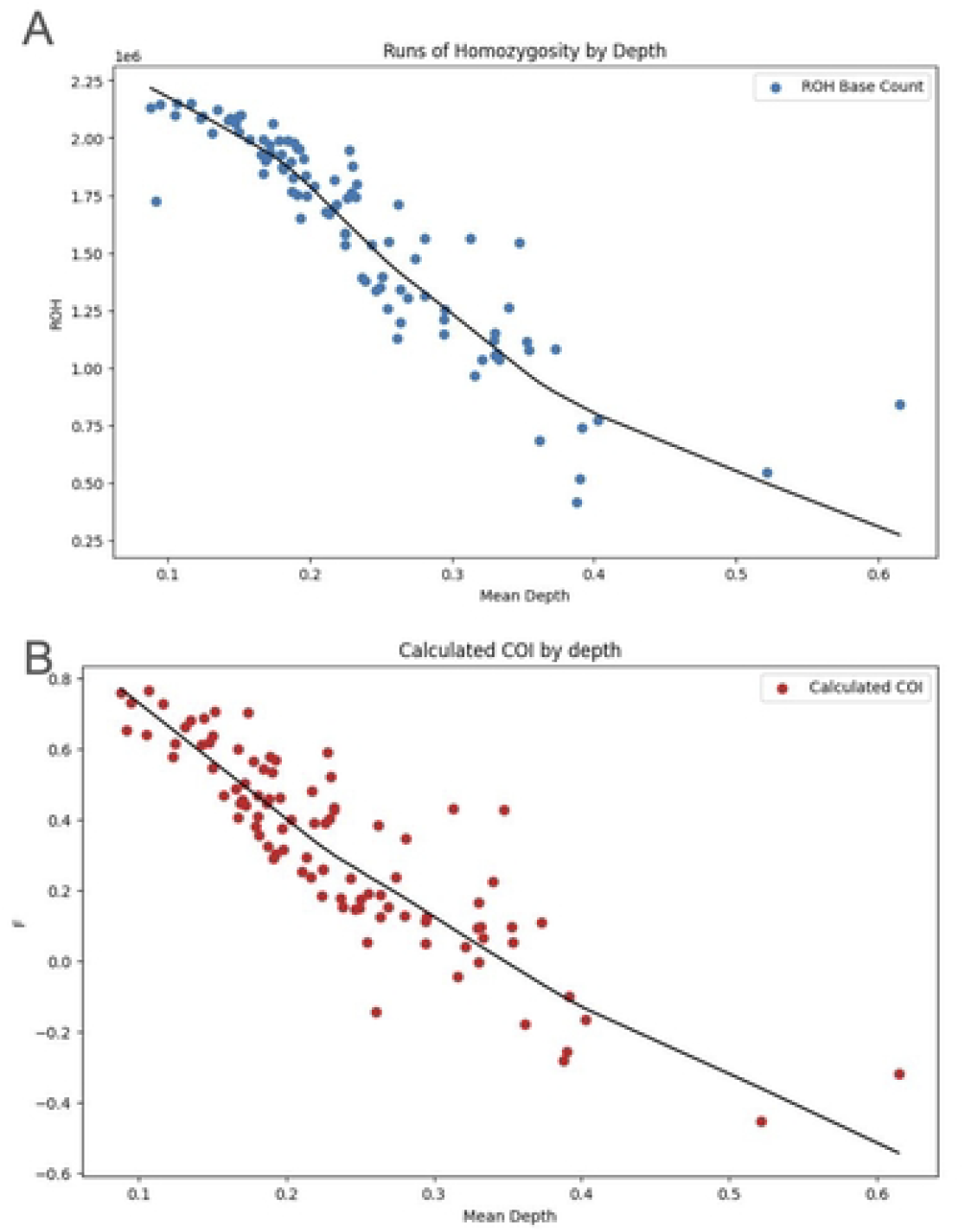
Inbreeding metrics and trends in respect of mean sequencing depth. (A) Runs of homozygosity and the depth of coverage show an inverse relationship. A LOESS regression line is shown in black. (B) Inbreeding coefficient and the depth of coverage show an inverse relationship. A LOESS regression line is shown in black.

To account for depth of coverage we implemented LOESS regression to compute the expected inbreeding values as a function of coverage. We found that using statsmodels.nonparametric.lowess() with frac=0.6 and it=4 for RoH values and frac=0.9 and it=5 for F values resulted in the best fit regression model (Fig 1). Residuals between the inbreeding value of one individual and the expected value for that coverage were then calculated by subtracting the actual inbreeding and RoH values and the predicted values of the model (Fig 2). These estimates of inbreeding represent the relative inbreeding of that individual in a population, and not the actual inbreeding values. An additional regression model fitted to the residuals show no significant correlation with variable sequencing depth, confirming that the coverage variability has been accounted for. We found that the inbreeding coefficient residuals and RoH residuals have a positive linear correlation with the coefficient of 0.834 and p-value of 4.8×10^-26^, indicating a general trend of increased inbreeding when the values of the two metrics increase (Fig 3).

**Fig 2.**
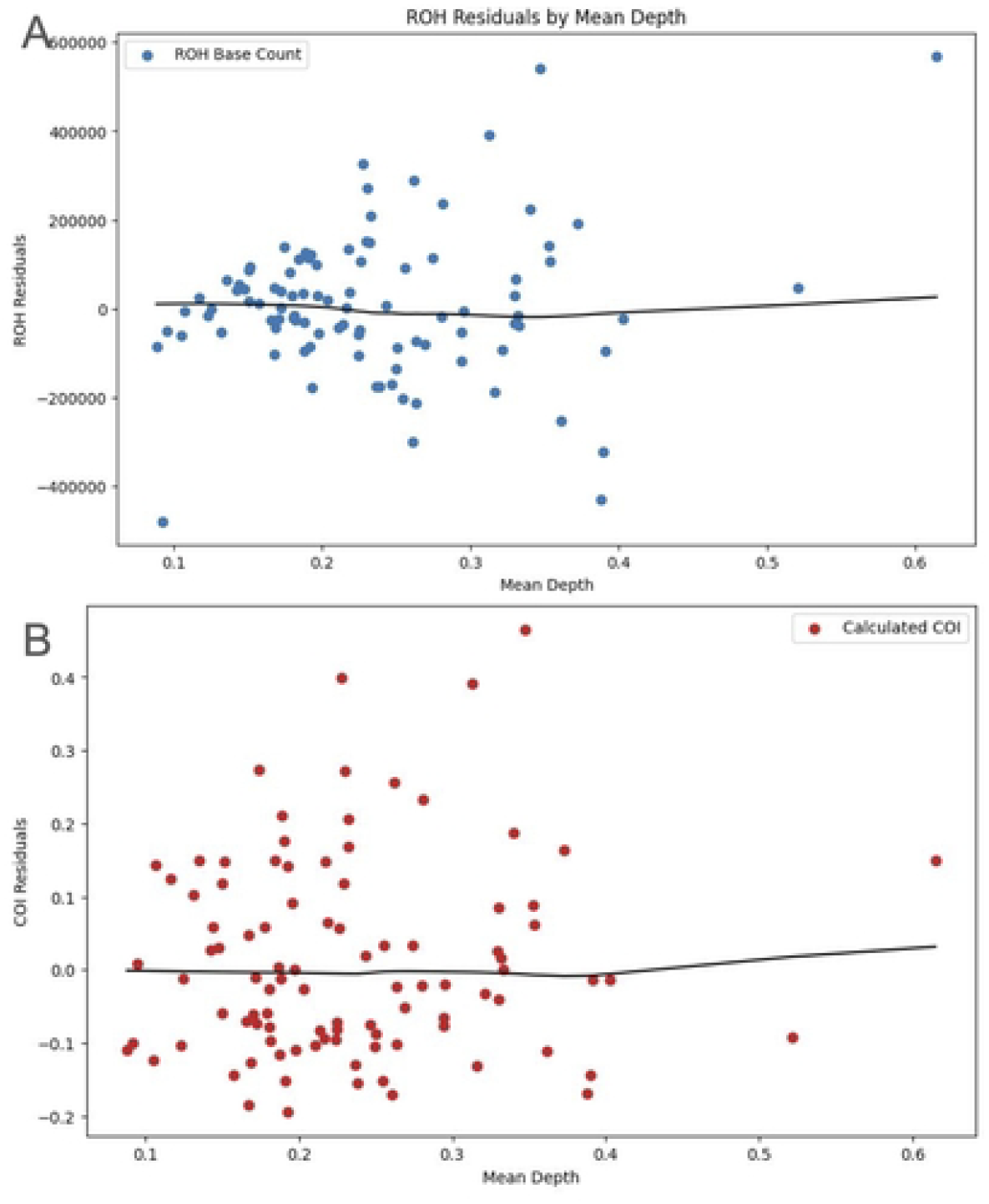
Residual calculation of inbreeding metrics to account for variable sequencing coverage. (A) Differences between individual RoH values and the expected values based on the LOESS model from Fig 1A versus mean sequencing depth. The LOESS fit of this data is shown with the black line. (B) Differences between individual F values and the expected values based on the LOESS model from Fig 1A versus mean sequencing depth. The LOESS fit of this data is shown with the black line.

**Fig 3.**
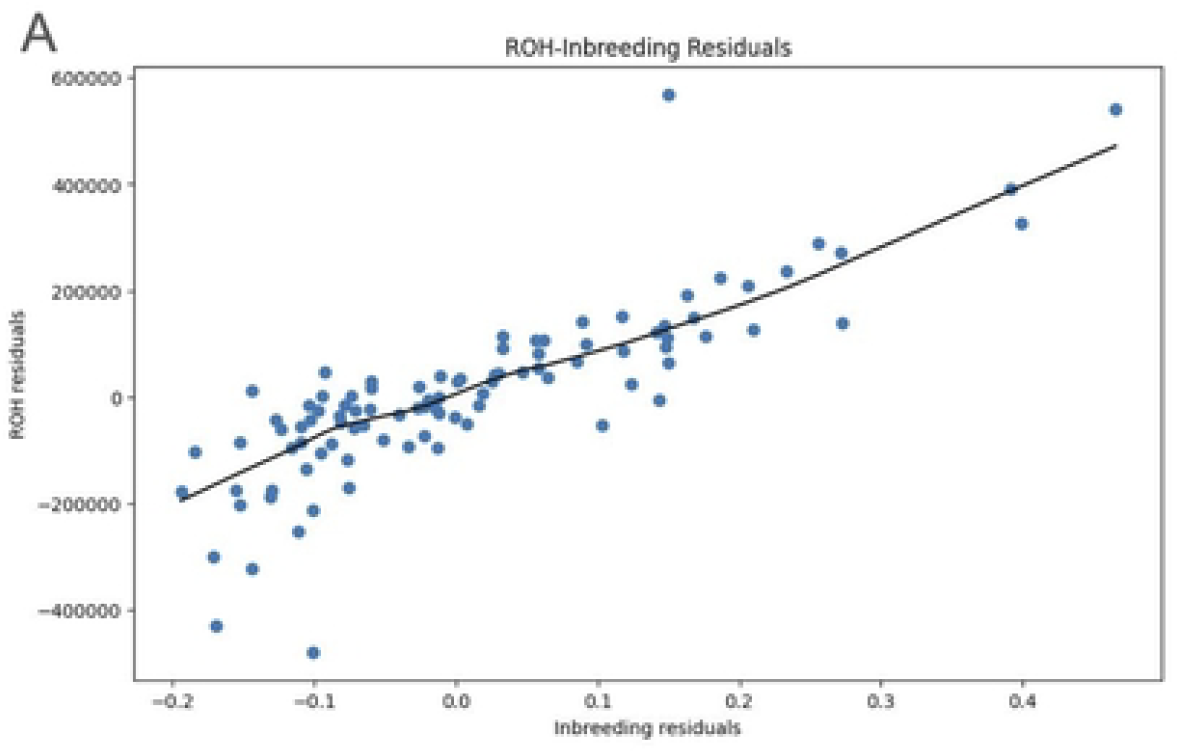
Correlation of inbreeding residuals. Residuals for both inbreeding metrics show a positive correlation, indicating a general trend of increased inbreeding when the value of two metrics increase.

To further confirm the relationship between inbreeding and coverage we generated a synthetic dataset by downsampling a high coverage WGS sample. The downsampled data exhibited a similar trend between inbreeding and sequencing depth, supporting the trend line between inbreeding and coverage that was used to calculate residuals (Fig 4).

**Fig 4.**
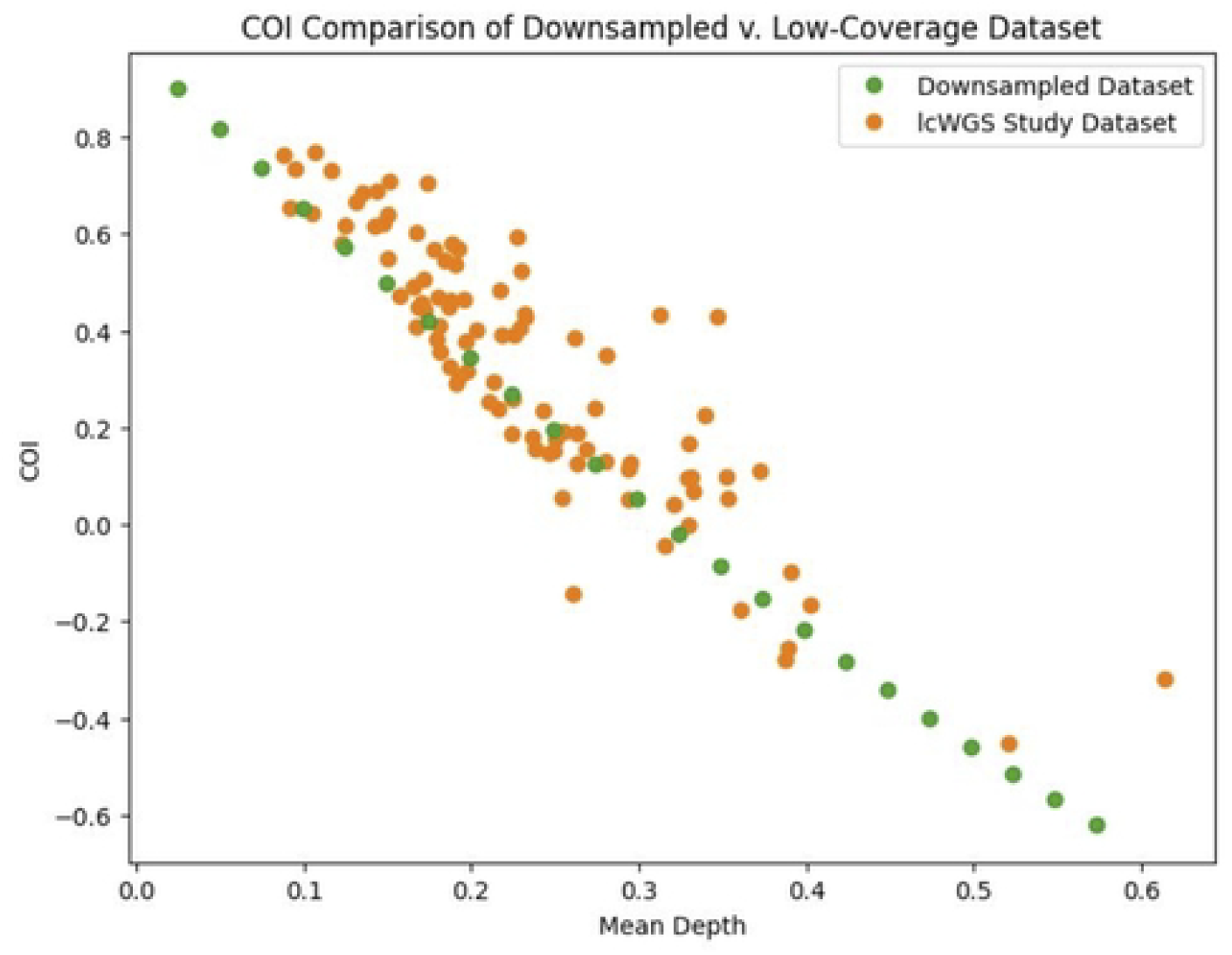
F Inbreeding versus coverage in synthetically downsampled dataset. F values for the ulcWGS dataset (orange) and the downsampled dataset (green) versus coverage.

### Inbreeding across pure and admixed dog breeds

Ranking of inbreeding across our low coverage samples was determined using the sum of ranks using RoH and F. Eight out of 10 dogs of the top 10 were reported as purebreds, compared to 8 out of 10 dogs of the lower 10 reported as mixed breeds (Fig 5). Furthermore, we found that breeds previously reported to exhibit higher levels of inbreeding, such as West Highland Terrier, Irish Wolfhound, Bloodhound, Airedale Terrier, Cavalier King Charles Spaniel, Rottweiler, Bullmastiff, and Bedlington Terrier, accounted for 12 of the top 20 most inbred pure breed individuals in our dataset [19]. Notably, out of the 11 breeds in our sample that fall within the top quartile of published inbreeding levels, 10 breeds were represented in our top 20 most inbred individuals, further supporting the consistency of our findings with established breed-level inbreeding patterns (Fig 6). Among the 11 breeds with lower values, four were in the bottom quartile of published inbreeding levels. More outbred breeds include the Australian Shepherd, Chihuahua, and Jack Russell Terrier. On average, purebreds had a mean RoH residual that was 98 kbp higher (Welch’s t-test; t = 2.69, p = 9.65×10^-3^) and an COI residual that was 0.99 higher (Welch’s t-test; t = 3.38, p = 1.34×10^-3^). Additionally, the overall inbreeding rankings were significantly higher for purebreds (Mann-Whitney U = 1598, p = 8.299×10^-6^).

**Fig 5.**
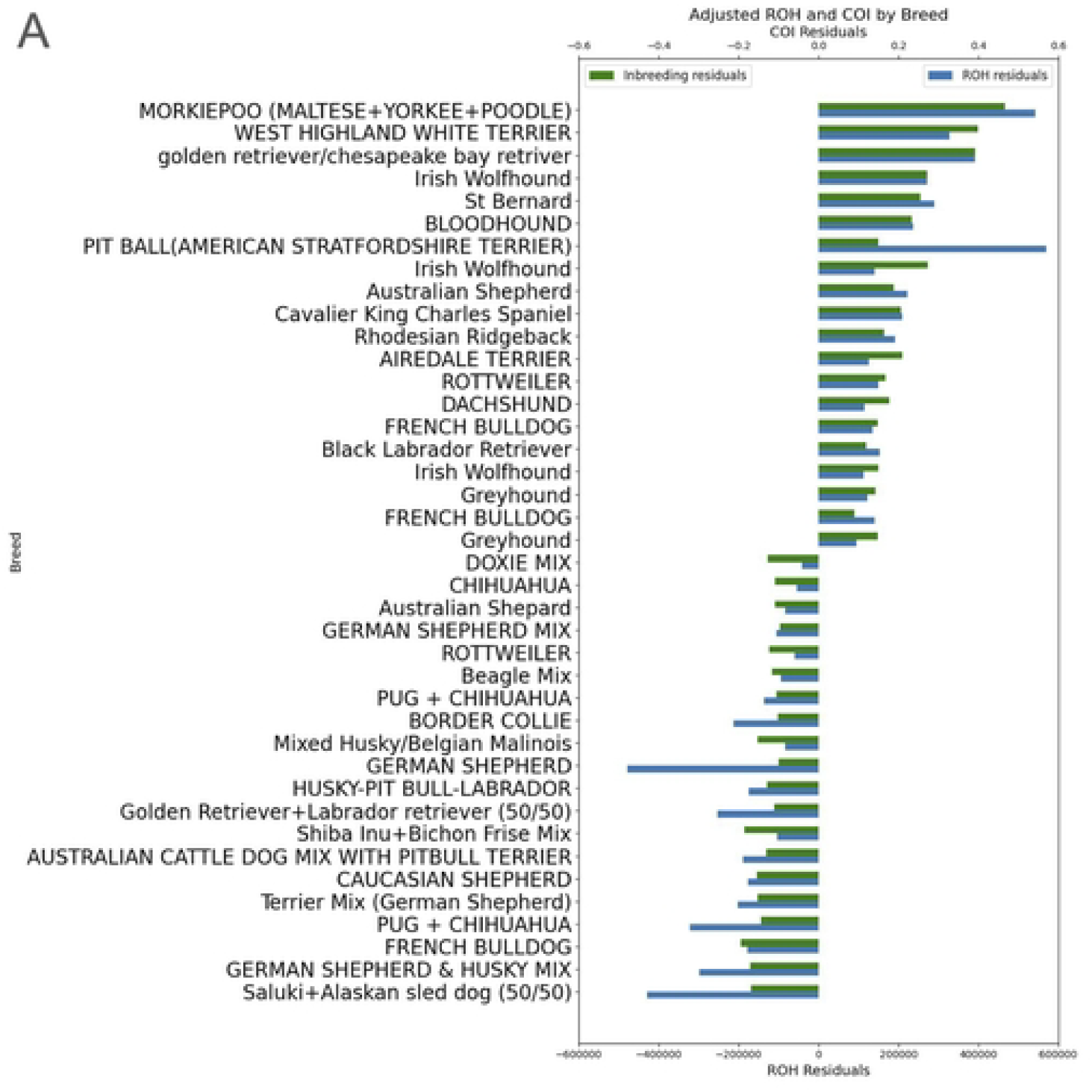
Adjusted inbreeding metrics of select individuals listed by breed. The graph shows adjusted F (green) and RoH (blue) of samples on the x-axis by breed on the y-axis. Each sample was listed by the average ranking, and the top and bottom 20 are shown.

**Fig 6.**
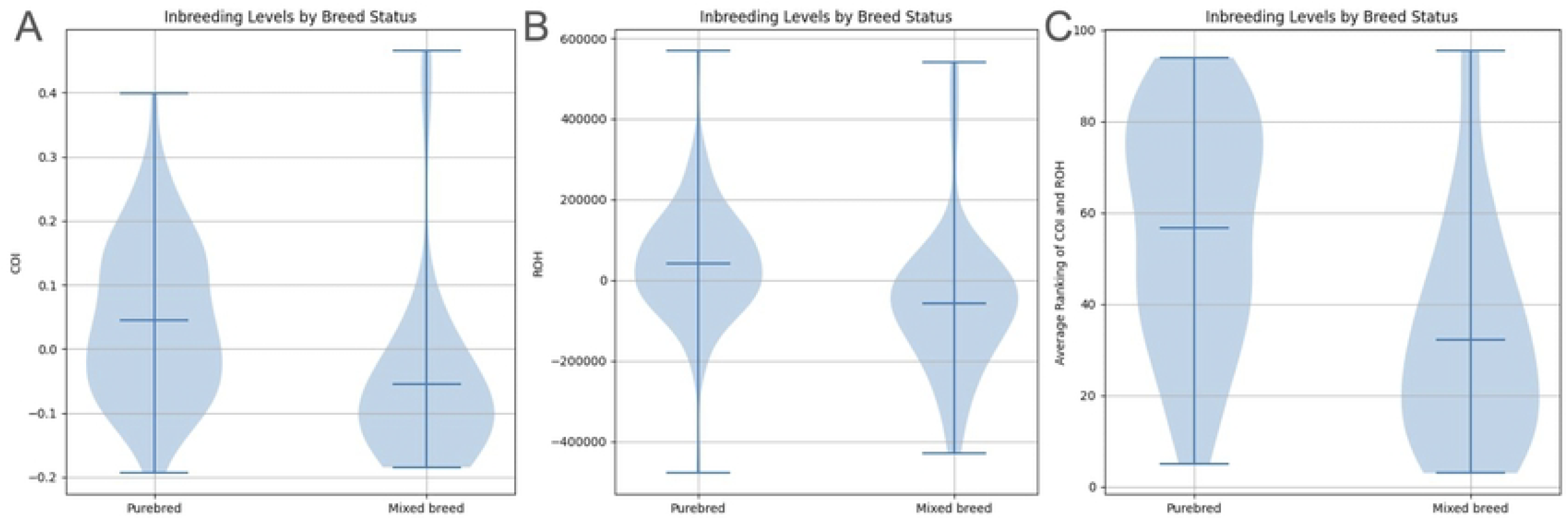
Inbreeding status by breed type. (A) Estimated F is shown by breed status. The line in the center of the violin plot shows the mean F value. (B) Total base count of RoH is shown by breed status. The line in the center of the violin plot shows the mean base count. (C) Inbreeding ranking is shown by breed status, determined through the sum of individual F and RoH rankings. The line in the center of the violin plot shows the mean ranking of each type of breed.

## Discussion

Our study demonstrates that ultra low-coverage genome sequencing with an average depth of 0.1x-0.6x can provide estimates of relative inbreeding when they are corrected for sequencing depth. Previously, most studies estimating inbreeding have relied on high-coverage WGS (>15x). Skepticism remained when lower depth data (∼7x) was used, and such efforts often overlooked biases introduced by variable coverage [16]. We demonstrate that even at sequencing depths well below 1x, inbreeding signals can be recovered from ulcWGS data when corrected for sequencing depth. Our results were supported by previous estimates of variation in inbreeding levels across breeds.

Using LOESS regression to model the changes in inbreeding metrics as sequencing depth changes, we have attempted to address concerns about systematic biases in previous studies that used low coverage WGS with varying sequencing depths. Our simulation using downsampled data closely aligns with the trend seen in real samples. We observe that for the RoH analysis, in accordance with previous studies reporting both underestimations and overestimation of RoH at medium coverage around 6x-9x [21], we found deviations at the extreme ends of our ulcWGS data from 0.1x-0.6x. RoH analysis at the lowest coverage of 0.1x underestimated the number of RoH fragments, while at the highest coverage of 0.6x overestimated the number of fragments.

As ulcWGS is considerably less resource-intensive than high-coverage WGS, it offers a more accessible genetic tool for research, conservation, agricultural, and commercial applications. We find in this study that ulcWGS can be reliable for inbreeding estimation provided that the estimated inbreeding metrics are corrected for sequencing depth, highlighting its future potential as a scalable tool.

Several limitations should be noted. First, breed identity in our dataset was self-reported, which introduces uncertainty in breed assignments and may contribute to classification errors between purebred and mixed-breed individuals. We did not have higher-coverage WGS data for the samples, therefore did not have an independent inbreeding estimate to compare to. Second, while our sample size (96 individuals) was sufficient to demonstrate proof-of-concept, it was not designed to provide breed-level population estimates of inbreeding. Larger sample sizes with independently verified breed designations will be necessary to expand upon these findings. In addition, rare and vulnerable breeds were not fully represented in our sample pool. Finally, while we had relative differences in inbreeding across individuals and groups, we did not have a way to calibrate these values against an independent measure of the coefficient of inbreeding. In summary, the inbreeding measures reported here do not represent quantitative measures of inbreeding but rather are relative indicators with respect to a population.

Further work could explore different measures of inbreeding coefficients like F_UNI_, F_GRM_, and other methods to capture a more comprehensive view of various inbreeding metrics to be used on ultra low-coverage WGS data. Finally, expanding the scope of the study to other species, especially endangered ones, would enable the assessment of the broader utility of ulcWGS in conservation genetic monitoring and management.

## Data Availability Statement

The data underlying this study are available in the NIH sequence read archive under the BioProject ID of PRJNA1259595. The source code for the modification of the data and figures can be found with this GitHub link.

https://github.com/JRyanKim/Estimation-of-Inbreeding-using-ulcWGS

Figures

Sample Reads:

SubmissionID: SUB14934465

BioProject ID: PRJNA1259595

Reference Population:

Supplementary Table 1: [modsixclust](https://docs.google.com/spreadsheets/d/1SZ82710jMjmkVHLD5xzyfvPLUYrRHpH1MfS8qkZAmpE/edit?usp=sharing)

https://dataview.ncbi.nlm.nih.gov/object/PRJNA1259595

## Supporting information

**S1 Fig. Residual comparison by breed**

**S2 Fig. Complete rank of inbreeding metrics by breed**

**S1 Table**.

